# The Premotor Language Area Encodes a Full Acoustic-to-semantic Speech Hierarchy

**DOI:** 10.64898/2026.07.01.735929

**Authors:** Sihang (Suna) Guo, Alexander G Huth

## Abstract

Classic neurobiological models of human speech and language have emphasized the dominant role of temporal lobe in speech perception, while premotor regions including the ventral premotor cortex (PMv) are situated at the level of articulatory processing. However, accumulating evidence from neuroimaging, clinical, and computational studies suggests that premotor cortex may contribute to speech processing beyond articulation. The precise extent and functional organization of these speech-related representations, however, remain unclear. In this study, we combined naturalistic speech perception with computational encoding models to characterize the organization of speech representations within PMv. We functionally localized a cortical region that encompasses previously described premotor speech areas, which we term the premotor language area (PML). Using acoustic, phonemic, semantic, and deep neural speech representations, we found that PML contains representations spanning the full speech-processing hierarchy, from low-level acoustic features to high-level semantic information. These representations are arranged along a smooth posterior-anterior gradient, with increasingly abstract speech representations emerging toward anterior PML. Moreover, this organizational gradient mirrors the canonical speech processing hierarchy in the temporal auditory regions. These findings challenge the traditional view of premotor cortex as primarily an acoustic-articulatory unit, and instead identify PML as a hierarchically organized speech-processing region that parallels the temporal auditory cortex. This provides a new framework for understanding the role of premotor cortex in speech perception.

## Introduction

Spoken language processing is one of the most complex motor–cognitive behaviors humans perform, requiring the seamless integration of sensory analysis, motor planning, and linguistic computation. Achieving these tasks requires coordination between multiple areas of the cerebral cortex. Traditional neurobiological models posit crucial roles for the superior temporal lobe (STL) in auditory and speech perception, as well as frontal lobe areas in motor planning during speech production, including the inferior frontal gyrus (IFG; Broca’s area) and the ventral premotor cortex (PMv) (Hickok & Poeppel, 2004).

Over the past two decades, the premotor cortex has received increasing attention for its dual contributions to both speech perception and production. Early functional localization of this area found local activations during both perception and production of syllables and phonemes (Wilson et al., 2004; Pulvermüller et al., 2006; Cheung et al., 2016). Activations here also increased during effortful processing of syllables that are difficult (Wong et al., 2008; Tremblay & Small, 2011), unfamiliar (Callan et al., 2004; Wilson & Iacoboni, 2006), or masked by noise (Callan et al., 2010; Du et al., 2014). Conversely, disruption of this region via lesion or ablation impairs perception of syllables (Meister et al., 2007; D’Ausilio et al., 2011; Sammler et al., 2015; Higashiyama et al., 2021). These results have been interpreted as evidence for theoretical frameworks such as the motor theory of speech perception, which proposes that speech production systems are recruited to facilitate perception, particularly under demanding conditions (Callan et al., 2010).

More recently, studies using naturalistic speech tasks further revealed that our target region exhibits cross-complexity selectivity spanning acoustic, articulatory, and semantic domains. Activity in this region can be well predicted by cochleograms, phoneme identities, and word embeddings (Heer et al., 2017), as well as contextual acoustic and semantic embeddings derived from large speech and language models (Antonello et al., 2023; Vaidya et al., 2022). A recent study on speech production also shows that anterior and posterior area 55b exhibit differential encoding for semantic and articulatory features (Wu et al., 2026). This broad representational profile is corroborated by meta-analyses in psycholinguistic and patient studies: alongside canonical speech processing areas including IFG and pSTG, the premotor area contains activation foci for phonological, lexical semantic, and sentence-level syntactic perception and production (Vigneau et al., 2006), as well as lesion foci accompanied by anarthria, phonological errors, semantic errors, and anomia (Lu et al., 2021; Tate et al., 2014). Together, these aggregated results indicate the crucial role of this region at multiple levels of speech processing across perception and production, in both natural and controlled experimental paradigms.

Despite this growing body of evidence, the functional organization of this cortical territory remains poorly understood. Localization of this area has mostly been done using controlled tasks designed to isolate specific levels of speech processing (e.g., phonemes, words, and sentences), yielding definitions of overlapping but non-identical portions of the PMv. Literature in different subfields has also referred to the general area separately as the superior ventral premotor cortex (sPMv) (Wilson et al., 2004), the middle precentral gyrus (midPrCG) (Silva et al., 2022), the dorsal precentral speech area (dPCSA) (Hickok et al., 2023), the middle frontal gyrus (MFG) language area (Fedorenko et al., 2024), and area 55b (Hopf, 1956; Glasser et al., 2016). While these regions are often treated as functionally equivalent, different definitions may in fact reflect multiple adjacent subregions selectively engaged by distinct stimulus features.

Consequently, several fundamental questions remain unresolved. What is the full functional extent of this cortical area in speech and language perception? Does this region encode only isolated linguistic representations, or does it also support transformations between acoustic, articulatory, and semantic levels of processing? Furthermore, how are these diverse representations spatially organized? Are the neural populations encoding different levels of speech features interspersed across the entire region, or aligned with some anatomical-functional axis?

In this study, we address these questions by examining neural responses during naturalistic speech perception using computational encoding models. We define a unified functional region of interest that encompasses earlier definitions, which we term the **premotor language area (PML)**. Natural speech and encoding models allow us to simultaneously map acoustic, phonemic, and semantic representations within the same subjects, and compare their anatomical distributions. We further compare representations derived from interpretable linguistic features with those obtained from large speech models that capture continuously varying speech structure. We test competing hypotheses about the functional organization within PML and show that PML exhibits a posterior–anterior hierarchical gradient, with increasingly complex speech representations encoded as the ROI extends from the primary motor cortex (M1) to the middle frontal gyrus (MFG). Using functional correlation, we further demonstrate that the functional gradient in PML mirrors the canonical speech processing hierarchy observed from primary auditory cortex (A1) to lateral and posterior superior temporal gyrus (STG). We also rule out the possibility that PML is part of the somato-cognitive action network (SCAN) (Gordon et al., 2023), despite its close proximity. Together, these findings establish PML as a hierarchically organized component of the speech-processing network and provide new insight into how premotor cortex contributes to the integration of perception, action, and language.

## Results

### Localizing Premotor Language Area (PML)

Previous studies have functionally identified subregions of PML by intersecting the areas identified by speech perception and production tasks, often using a single level of speech representation, such as phonemes. We believe that this approach, while specific, misses the full picture of speech and language representations in the area.

Here, we functionally defined PML using more naturalistic perception and production tasks. For perception, participants listened to mixed audio clips that included music, natural sounds, and natural speech. This stimulus set covers a broader range than just speech, allowing inclusion of more acoustically selective regions as well. For production, we used an imagined speech task, where participants generated internal speech without voicing or mouth movements. This approach decouples speech production from motor movements, resulting in a cleaner localization of higher-level speech production.

We then defined PML for each subject as the union of voxels that were either strongly activated by the audition task or the imagined speech task (**Figure 1A**). While the audition task mainly activated posterior PML, the imagined speech task mainly activated the anterior PML, with overlapping activations around the anterior border of M1M, the primary mouth motor cortex (**Figure 1B**). Consistent with the general location in previous studies, PML is found at the junction of the middle frontal gyrus (MFG) and the precentral gyrus (preCG), spanning portions of the primary motor (M1) and premotor cortex (PM) (Brodmann areas 4/6). In all subjects, posterior PML is located on the crown of preCG, partially overlapping with the primary mouth motor cortex (M1M). Anterior PML is more variable: for some subjects, it is on the gyrus connecting preCG and MFG; for others, it passes through the anterior extension of the precentral sulcus. Inter-hemispheric differences were observed as well. One subject exhibited the rare organization of dual PML on the same hemisphere, where two unconnected regions respond to both speech perception and production (Glasser et al., 2016). This subject was excluded from other analyses, but is included in individual maps (**Figure S1**) to illustrate the diversity of PML functional anatomy.

**Figure 1.**
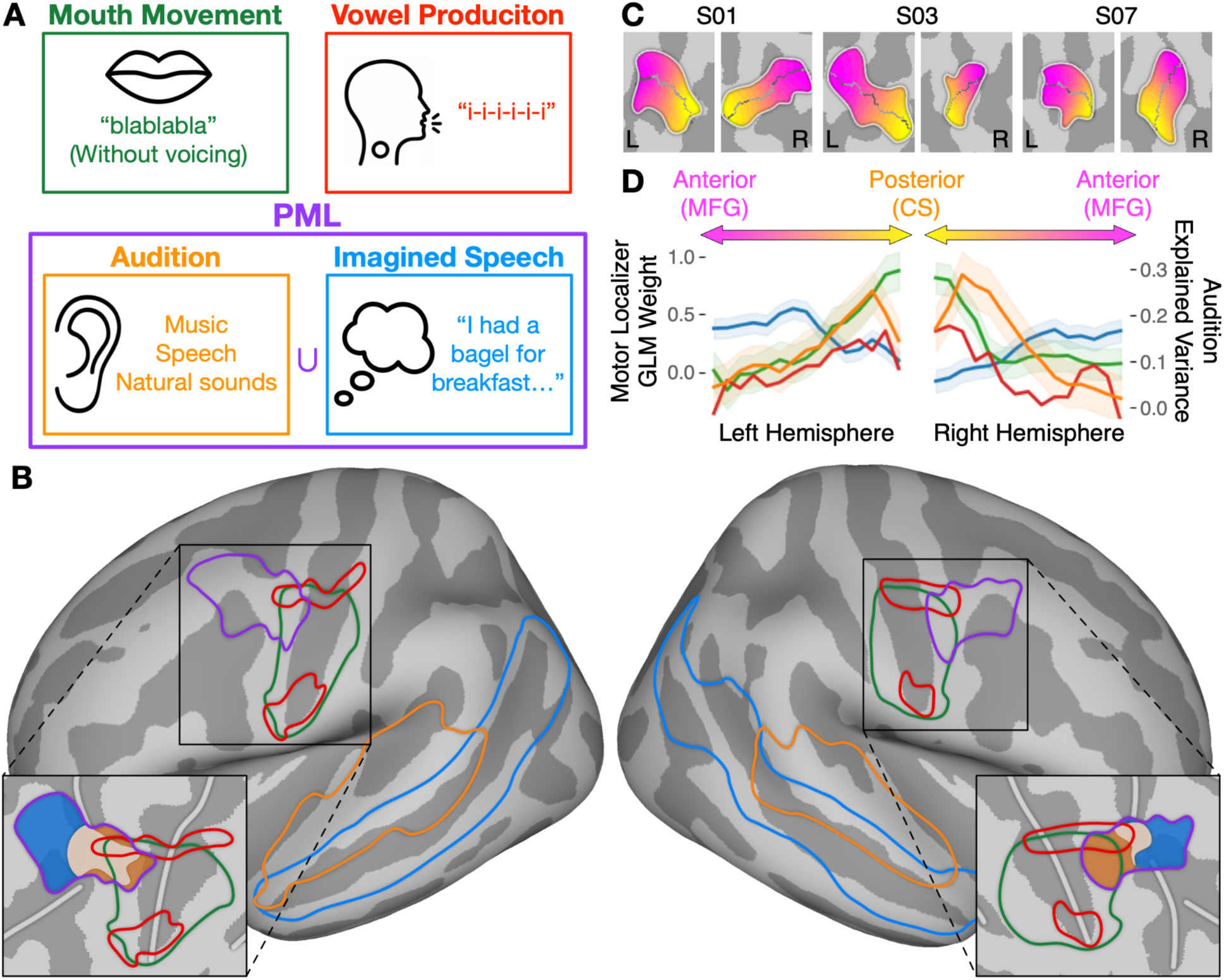
The premotor language area (PML) and other functional areas localized in the (pre)motor region. All ROIs are hand-drawn from contrast maps, not based on a prespecified significance threshold. **(A)** Four localizer tasks were used to functionally define PML (purple), primary mouth motor and somatosensory area (S/M1M; green), and laryngeal motor cortex (LMC; red). **(B)** Aggregated localization for each ROI across 7 subjects, outlined with corresponding colors from tasks in (A). Overlapping activation ROIs for audition and imagined speech appear in temporal lobe and as shaded areas within PML in inset map; overlap in PML is shaded in white. **(C)** PML is divided into bins along the posterior-anterior (yellow-magenta) axis for three sample subjects. **(D)** Localizer measures for each bin across the AP axis, averaged across subjects. Anterior PML responds more selectively to imagined speech, while posterior PML shows stronger responses to audition, mouth movement, and vowel production. The shaded area indicates standard error. The vowel production task was measured in one subject only, so no error measure is plotted. MFG: middle frontal gyrus; CS: central sulcus.

Intracranial EEG studies suggest that PML is closely tied to or colocalized with the dorsal laryngeal motor cortex (dLMC; Silva et al., 2022), which is also activated during both speech perception and production (Cheung et al., 2016; Dichter et al., 2018). We tested this possibility by localizing dLMC in one subject and comparing its location to PML. As shown in **Figure 1B**, voxels consistently activated by laryngeal movements (vowel production task) tend to lie more posterior into the central sulcus across motor and somatosensory cortex, whereas PML is located on the crown and anterior of the preCG. There is partial overlap between dLMC and posterior PML, suggesting that PML uses laryngeal signals for pitch and voicing processing. This may support the conversion between articulatory and phonological representations as suggested by previous studies (Fedorenko et al., 2010).

Our localization also indicates that PML is elongated, spanning M1 and MFG. Since the anatomical location bridges lower-level primary sensorimotor regions and higher-level associative regions, we hypothesized that the speech features in PML are anatomically organized along the anterior-posterior (AP) axis. To test this hypothesis, we divided PML into 15 bins along the AP axis (**Figure 1C**). We then averaged measurements and modeling results within each bin to show their spatial organization. For example, in **Figure 1D**, we show the measurements for all localizer tasks across this AP axis: posterior PML shows stronger selectivity for audition, mouth movement, and vowel production tasks, while anterior PML is more selective for imagined speech.

### Establishing the PML Speech Processing Gradient

Previous literature suggests that representations for many levels of speech–from pitch, acoustic, phonological, to semantic–can be found in or around PML. However, most prior studies have relied on tightly controlled stimuli that isolate a single level of speech representation. While effective for detecting task-specific responses, such paradigms make it difficult, and often inaccurate, to integrate results across studies and subjects to discern any spatial organization that might be present across these representations.

To overcome these limitations, we used naturalistic speech stimuli that engage all levels of the speech processing simultaneously in the same subjects. Six participants from the localization tasks listened to over six hours of narrative speech from the Moth Radio Hour while whole-brain blood-oxygen-level-dependent (BOLD) signals were recorded using fMRI.

To understand how selectively each voxel responds to different speech representations during naturalistic listening, we obtained speech features at various levels of representation.

First, we extracted speech features using self-supervised speech models (S3Ms)–artificial neural networks trained on large-scale audio corpora to learn hierarchical speech representations without explicit linguistic supervision (Mohamed et al., 2022). Specifically, we used the large variant of WavLM, a transformer-based S3M that is the state-of-the-art model on a battery of speech and language tasks, as well as brain modeling (Chen et al., 2022; Antonello et al., 2023). Prior work has demonstrated that representations in WavLM increase in complexity across layers, with early layers representing low-level acoustic features, intermediate layers representing phonetic information, and upper layers representing high-level lexical information as well as traces of phonetic and acoustic features through residual connections (Pasad et al., 2023; Vaidya et al., 2022). This continuous hierarchy makes WavLM an ideal model for testing whether a functional gradient exists within PML for speech perception. We passed the stimulus audio through WavLM and extracted hidden states from each layer as speech features across the hierarchy.

We then used voxel-wise encoding models to relate these speech features to BOLD responses. For each voxel, we used regularized linear regression to estimate how strongly its activity was modulated by features extracted from each WavLM layer. Model performance can be evaluated by predicting BOLD responses to held-out stimuli and computing the correlation between the predicted and observed responses (**Figure 2A**). This provides a measure for the fraction of the response variance explained by the features in each WavLM layer. Since WavLM layers show increasingly complex speech representations towards upper layers, the prediction performance of different layers should reflect the features encoded in each voxel within PML. For example, the encoding performance of lower layers reflects the encoding of acoustic features, while the performance of the upper layers reflects lexical and semantic features as well as acoustic features.

**Figure 2.**
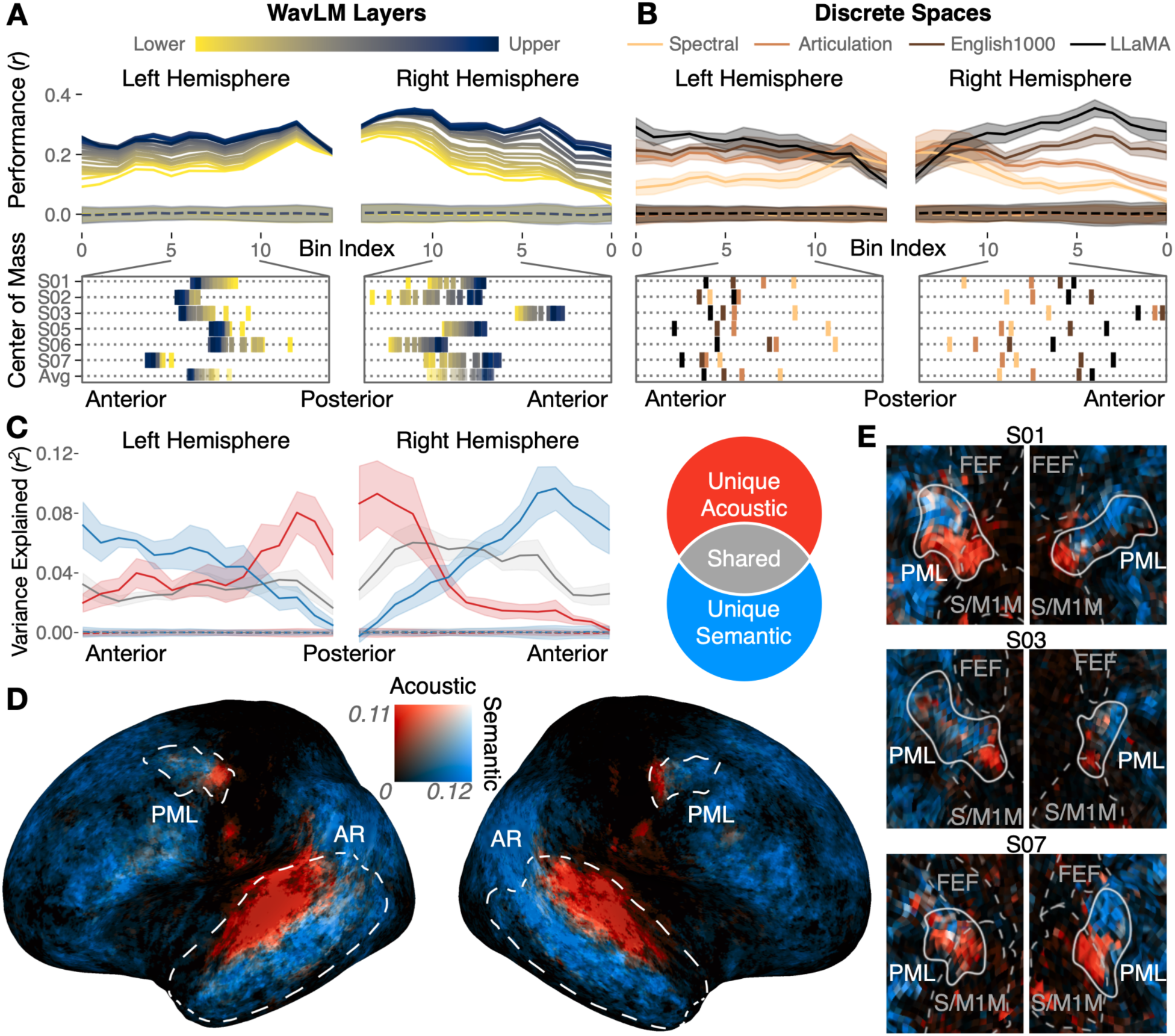
Speech representations in PML follow a functional gradient that is aligned with the anterior-posterior axis. **(A** & **B)** Top: Encoding performance of WavLM layers (A) and other discrete speech features (B) shows that acoustic features (lower WavLM layers, spectral, and articulation) well model posterior PML only, while anterior PML is well modeled by semantic features (upper WavLM layers, word embedding, and contextual embedding) only. All models predicted PML bin performances significantly better than chance (p < 0. 001, permutation test, FDR corrected), except for spectral and WavLM layer 0 in the two most anterior bins in the right hemisphere. Bottom: center of mass across bins for each feature space shows a gradual increase in model performance towards anterior bins for higher-level features, indicating a smooth functional gradient for speech representations in PML. **(C)** To disentangle acoustic and semantic representations in PML, we used a nested regression approach and partitioned the variance explained by WavLM layer 9 alone (acoustic, red), LLaMA layer 18 alone (semantic, blue), and shared by both feature spaces (grey). LLaMA layer 18 (semantic, blue) explains little unique variance in posterior PML but high variance in anterior PML, while WavLM layer 9 (acoustic, red) shows the opposite trend. This suggests that posterior PML encodes mainly acoustic information and anterior semantics. Both feature spaces show a smooth transition of unique variance explained in the middle bins, indicating a gradient of speech representations within PML. All models explain significant amount of variance in all bins except for (i) unique acoustic in the most anterior bin, and (ii) unique semantic in the most posterior bin, in the right hemisphere (p < 0. 001, FDR corrected, permutation test). **(D)** Unique variance explained by each model, averaged across six subjects. Distinct areas in PML and temporal auditory region (AR) are well explained by acoustic features alone (red) or semantic features alone (blue) (p < 0. 001, FDR corrected, permutation test). PML shows a posterior-anterior gradient for acoustic-semantic features, mirroring that in STG and MTG. **(E)** Unique variance explained by each model for three example subjects. FEF: frontal eye fields; S/M1M: primary somatosensory/motor mouth area.

The prediction performance in **Figure 2A** shows a continuously evolving representation gradient in PML. Lower WavLM layers only perform well in posterior PML, suggesting that selectivity for low-level speech features is concentrated there. In contrast, anterior PML is much better predicted by upper WavLM layers than lower layers, indicating that anterior PML encodes higher levels of speech representation. Upper WavLM layers show better prediction performance in all bins including posterior PML, reflecting the effect of residual connections between WavLM layers, which will be disentangled in the next analysis. To quantify the overall performance profile for each layer, we calculated the center of mass (CoM) of layer performance across all bins. Lower **Figure 2A** shows that the CoMs of upper WavLM layers shift increasingly anterior in both hemispheres, reflecting more complex speech representations in anterior PML.

Besides continuous representations from large speech models, we also included other commonly used speech features for reference, including spectral features, articulation (Mesgarani et al., 2014), word embeddings (English1000; Huth et al., 2016), and contextual embeddings (LLaMA 33B; Touvron et al., 2023). These are discrete levels of speech processing that align with linguistic definitions, whose cortical selectivity ranges from early auditory processing areas to broad associative regions (Antonello et al., 2023; Heer et al., 2017; Huth et al., 2016; Vaidya et al., 2022). Similar to WavLM features, chunks of stimulus audio were passed through each model and produced feature vectors for modeling cortical speech representations. **Figure 2B** shows encoding performance patterns for these features that tells a similar story to the WavLM layers: from lower to higher level speech features, the best predicted bins within PML gradually shift from posterior to anterior. These results confirm that not only does PML encode all levels of speech features that can be found elsewhere in the brain, but it also organizes the features in a meaningful pattern following the anatomical orientation.

The results above show a posterior-low-level, anterior-high-level functional gradient within PML. However, raw model performance in posterior bins appears to be similar across feature spaces. Does this indicate a mixed representation of acoustic and semantic features in posterior PML, with diverging selectivity towards the anterior end? Or could this be an artifact of residual connections in WavLM layers? Transformer-based models are made powerful in part by residual connections, which allow features to be passed from lower to upper layers. Therefore, once present, lower-level representations remain throughout the model, exhibiting a mixture of acoustic and lexical representations in upper layers of WavLM, which also achieve the best brain encoding performance (Antonello et al., 2023). The presence of such mixed representation could explain the similar performance across all layers in posterior bins, but these models alone do not account for the tuning properties of posterior PML.

To fully understand the structure of the functional gradient, we used variance partitioning to disentangle acoustic and semantic representations in PML. Variance partitioning uses nested regressions and set theory to estimate the unique contribution of each feature space to BOLD responses. First, we chose WavLM layer 9 and LLaMA (33B) layer 18 as our acoustic and semantic features, respectively. WavLM layer 9 is known for its performance on downstream acoustic tasks (Cho et al., 2024) and encoding performance in the human auditory cortex (Vaidya et al., 2022). LLaMA is a large language model trained for next-word prediction with word-level tokenization, hence it should mainly encode semantic features at or above the lexical level. Layer 18 of LLaMA showed the highest whole-brain modeling performance for natural language perception in a separate study (Antonello et al., 2023). To minimize shared low-level variance, we regressed out word rate from both feature spaces, such that the unique contribution of each feature space is not obscured. Using highly predictive features in both domains, variance partitioning should tell us whether posterior PML is a mixture of acoustic and semantic representations.

By fitting encoding models with features from WavLM layer 9 alone, LLaMA layer 18 alone, and the two combined, we can estimate the variance ( ^2^) uniquely explained by each feature set and the shared component. As shown in **Figures 2C** and **2D**, acoustic features (WavLM layer 9) uniquely explain the most variance in posterior PML. Meanwhile, semantic features (LLaMA layer 18) uniquely explain little, if any, variance in posterior PML, but much more in anterior PML. Both feature spaces showed a gradual change in unique ^2^ along the AP axis, but in opposite directions, demonstrating evolving speech representations within PML. Together, these results show that PML follows a posterior-acoustic, anterior-semantic functional gradient.

### The PML Functional Gradient Mirrors That of Temporal Auditory Regions

The canonical cortical substrate for speech perception is the temporal auditory regions (AR), where neural populations represent speech information across multiple levels of abstraction, from spectrotemporal acoustic structure to lexical and semantic content. Our results reveal that PML exhibits a remarkably similar organizational principle. Variance partitioning analyses showed that posterior PML preferentially encodes low-level acoustic information, analogous to medial auditory regions, whereas anterior PML preferentially encodes semantic information, resembling more lateral and anterior temporal regions. The presence of a complete acoustic-to-semantic representational gradient within this localized premotor territory raises a fundamental question: how is the functional organization of PML related to the canonical speech-processing hierarchy in the temporal lobe?

To answer this question, we computed the task-state functional connectivity (tsFC) between PML voxels and all cortical voxels, using voxel activations during over 6 hours of natural speech perception. For each cortical voxel, we calculated its average tsFC with all voxels in each PML bin. This allows us to examine how the functional gradient identified above interacts with other regions of the cortex, especially STG. For each cortical voxel, we then calculated the center of mass (CoM) for the average tsFCs across the bins, which indicates where along the PML A-P axis this voxel is more correlated (**Figure 3A**).

**Figure 3.**
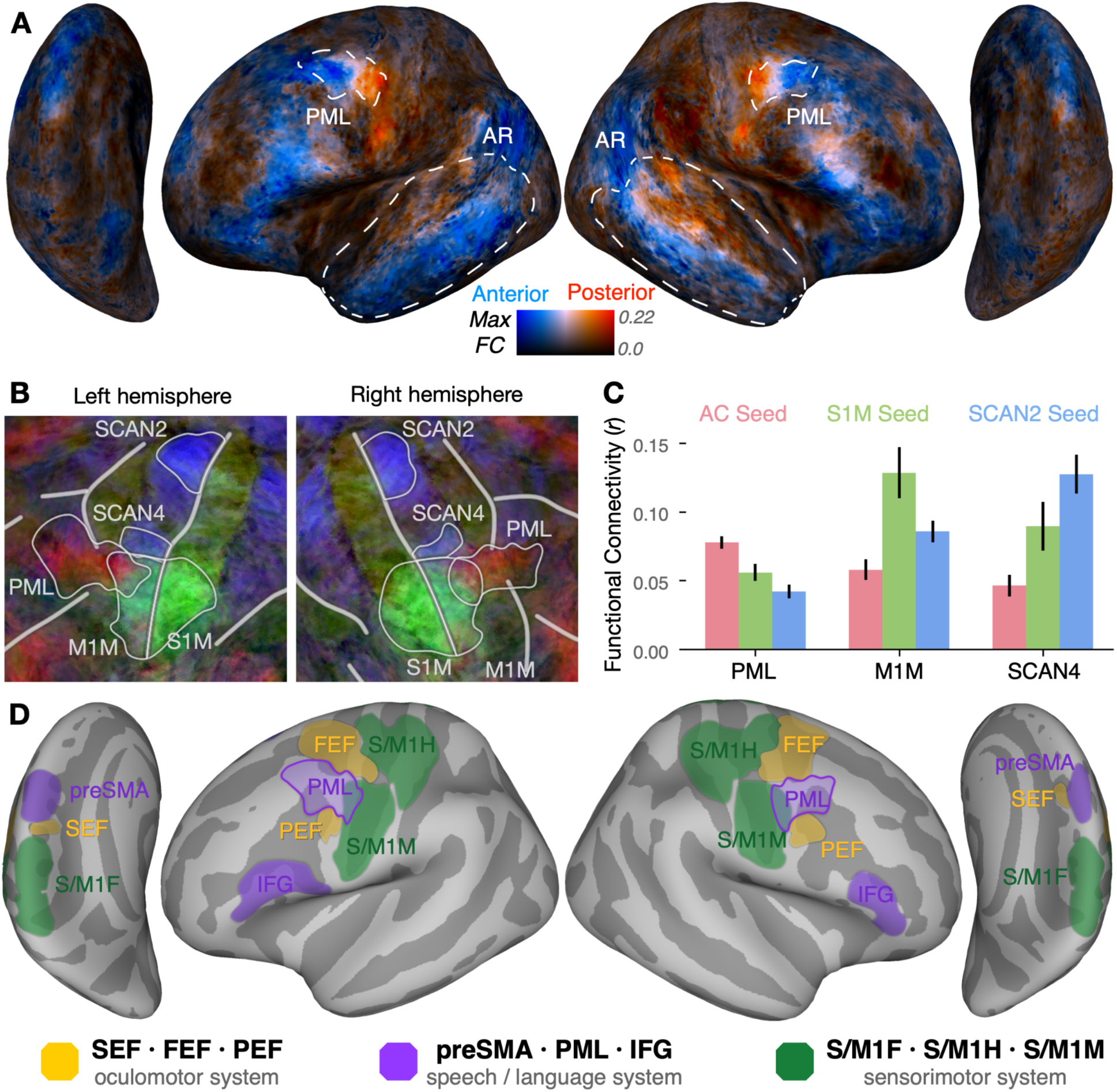
PML among other functional networks in the greater cortical landscape. **(A)** tsFC strength (hue) and preference to anterior (blue) or posterior (red) PML, for all cortical voxels. The PML functional gradient is correlated with the known speech processing pathway in temporal AR. Bin preference is calculated as the center of mass (CoM) for the average tsFC in each bin; strength is calculated as the max tsFC across bins. Each hemisphere shows the measure seeded from the ipsilateral PML, averaged across 6 subjects. **(B & C)** tsFC seeded from AC (red), S1M (green), and SCAN2 (blue), is the strongest in PML, M1M, and SCAN4, respectively (all FDR corrected p < 0. 05). Data is averaged across 6 subjects; error bars show standard error. **(D)** The primary motor (green), oculomotor (yellow), and language (purple) networks in the (pre)motor cortex, forming a medial-lateral-inferior architectural motif in the frontal cortex.

Posterior PML showed stronger functional coupling with auditory cortical regions, whereas progressively anterior portions of PML were preferentially correlated with middle temporal regions associated with higher-order linguistic processing. The resulting connectivity pattern formed a smooth gradient spanning temporal and premotor cortex, indicating that the representational hierarchy identified within PML is systematically aligned with the canonical hierarchy in auditory cortex.

The existence of this mirrored organization suggests that PML may play a broader role in speech perception than is previously assumed. Classical models have largely emphasized premotor cortex as a substrate for phonetic-articulatory transformations; although previous studies have investigated the overall functional connectivity between a subregion of PML and AR (Wilson & Iacoboni, 2006; Venezia et al., 2021), none have recognized its full range of speech representations. In contrast, our results reveal an acoustic-to-semantic representational hierarchy within a premotor region that overlaps with the articulator-related motor cortex.

Although, to our knowledge, PML does not receive direct auditory input from the medial geniculate nucleus, tract tracing in nonhuman primates has demonstrated projections from posterior auditory belt to anterior PM (Romanski et al., 1999), providing a potential pathway through which auditory information could reach premotor cortex early in processing (Hamilton et al., 2018). Such connectivity raises the possibility that speech representations evolve in parallel across temporal and premotor systems, giving rise to two anatomically distinct yet functionally aligned representational hierarchies.

### Situating PML in the (pre)motor networks

The premotor and motor cortex contain several large-scale functional networks that integrate information across sensory, cognitive, and motor domains. One prominent example is the somato-cognitive action network (SCAN), which has been proposed to support action planning and the cognitive control of coordinated movements (Gordon et al., 2023). SCAN occupies the inter-effector regions of motor and premotor cortex and consists of multiple functionally coupled subregions. Notably, SCAN4, located between the primary mouth and hand motor representations (M1M and M1H), coincides with our localization of PML. Given this spatial proximity and the established role of premotor cortex in speech motor functions, an important question is whether PML constitutes a speech-related component of SCAN or instead belongs to a distinct functional network.

To address this question, we compared the tsFC profiles of PML and SCAN. Direct comparisons between neighboring cortical regions can be confounded by spatial correlations arising from MRI signal smoothness and local vascular effects. We mitigate such confounds by using distant seed regions with established functional associations. Specifically, we seeded tsFC from auditory cortex (AC), which exhibits strong connectivity with PML (**Figure 3A**); primary mouth somatosensory cortex (S1M), which is strongly coupled to M1M; and SCAN2, the superior inter-effector region between the foot and hand representations (M1F, M1H). If PML belongs to SCAN, its connectivity profile should overlap substantially with that of SCAN4 and other SCAN regions. Conversely, distinct network membership should produce separable connectivity patterns.

The resulting connectivity maps revealed a clear dissociation (**Figure 3B-C**). SCAN2 showed strong connectivity with SCAN4 (blue), S1M was preferentially connected to M1M (green), and AC was preferentially connected to PML (red). Two-way ANOVA revealed a significant interaction between seed and target regions (*F*(4, 20) = 33. 97, *p* < 0. 001).

Paired t-tests comparing each ROI pair with non-pair FC showed that PML, M1M, and SCAN4 are most strongly connected with AC, S1M, and SCAN2, respectively (all FDR corrected *p* < 0. 05). Although the auditory and somatosensory connectivity maps are overlapped near posterior PML and superior M1M, connectivity associated with SCAN2 remained largely segregated from both networks. These findings indicate that PML is functionally distinct from SCAN despite their close anatomical proximity. Instead, the tsFC pattern between PML and AC (**Figure 3A**) closely matches the language network (Fedorenko et al., 2010), including signature regions such as pIFG, pSTG, as well as a premotor area around PML. Together, these results suggest that PML is part of the speech-language network rather than SCAN, establishing a functional dissociation between speech-related and action-control systems within the premotor cortex.

The dissociation between PML and SCAN also places PML within a broader organizational framework of the frontal motor system. Several functional systems in the frontal cortex are distributed across a set of anatomically separated but functionally related regions spanning the supplementary motor cortex and premotor cortex (**Figure 3D**). For example, the oculomotor system comprises the supplementary eye fields (SEF), frontal eye fields (FEF), and premotor eye fields (PEF), occupying medial, lateral, and inferior frontal territories, respectively. Intriguingly, speech and language regions exhibit a similar large-scale arrangement, with preSMA (Hertrich et al., 2016; Levy et al., 2025), PML, and IFG forming corresponding medial, lateral, and inferior frontal components. Although the precise functional roles of these regions remain to be established, this parallel organization suggests that speech-related processing may be embedded within a broader architectural motif of frontal cortex, in which specialized cognitive-motor systems are implemented across matched supplementary and premotor nodes.

## Discussion

In this paper, we demonstrated that naturalistic speech stimuli combined with voxel-wise modeling can reveal fine-grained functional organization in cortical regions that are difficult to characterize using traditional controlled paradigms. We identified a small premotor region, which we term the premotor language area (PML), that contains representations spanning the speech processing hierarchy. By modeling responses with interpretable linguistic features and self-supervised speech representations, we found that PML exhibits a posterior–anterior functional gradient, with progressively higher-order representations extending from acoustic to semantic levels. Moreover, this functional gradient mirrors that in the canonical temporal auditory regions. We further demonstrate that, despite its proximity to the somato-cognitive action network (SCAN), PML is functionally dissociable from SCAN, but is preferentially connected with the language network. Together, these findings establish PML as a hierarchically organized speech-processing region within the premotor cortex and reveal an unexpected correspondence between premotor and temporal-lobe representations of speech.

Although the present study examined speech perception alone, the results raise important questions regarding the broader cortical circuitry supporting speech and language. The existence of a complete acoustic-to-semantic gradient within PML suggests that speech representations are distributed across multiple cortical systems rather than confined to the temporal lobe. This observation raises several unresolved questions: At what stage of processing is information relayed from temporal auditory regions to PML? Which aspects of the observed hierarchy are inherited from upstream auditory representations, and which arise through local computations within PML? Furthermore, given the established involvement of PML in both speech perception and production, how does information flow through this circuit differently across behavioral contexts? Anatomical studies in both nonhuman primates (Romanski et al., 1999; Schmahmann et al., 2007) and humans (Axer et al., 2013; Dick et al., 2014; Shekari & Nozari, 2023) have identified structural connections between posterior superior temporal cortex and premotor regions near PML via the superior longitudinal fasciculus and arcuate fasciculus. However, the currently known anatomical pathways remain insufficient to explain the detailed functional gradient observed here. Intriguingly, among the three language-selective regions distributed across supplementary and premotor cortex (preSMA, PML, and IFG; **Figure 3D**), preSMA and IFG are linked by the frontal aslant tract, whereas no major white-matter pathway has yet been identified that places PML within this framework. This gap is particularly notable given that all three areas have similar functional profiles: ample studies show that all three language areas are involved in both speech perception and production, and all three areas contain representations spanning multiple levels of the speech processing hierarchy, similar to what we showed for PML in this paper (Ardila, 2020; Hertrich et al., 2016; Price, 2012). The broad range of speech features found in these perception-production interfaces suggests that key aspects of the general speech processing circuits remain to be understood.

Via task state functional correlation, we found that PML is dissociated from the SCAN. A recent study, however, suggested that area 55b, or posterior PML, is functionally connected to SCAN during speech production tasks aggregated across subjects (Wu et al., 2026). One possibility for this discrepancy is that voxel-wise modeling in individual subjects provides the precision required to discern area boundaries like this. Another possibility is that the difference in localization results from absence of speech-related movements in our study, which we specifically avoided in the localizer tasks. However, speech related activations are often attributed to effector regions such as LMC and M1M, while SCAN is localized specifically in the intereffector areas. Therefore, if the responses observed in PML primarily reflected overt or covert motor activity, one would expect the activations to extend posteriorly into the central sulcus, rather than superiorly into SCAN4. Nevertheless, SCAN may participate in speech production in a more nuanced way: previous studies have reported stronger activation in the trunk motor cortex and another PMv region–both around the two superior SCAN areas–during voiced speech compared with whispering, suggesting respiratory control and whole body coordination during vocalization (Correia et al., 2020; Paus et al., 1996). Thus, whereas PML appears to participate directly in speech and language processing, SCAN may contribute to the broader sensorimotor coordination required for speech production. Clarifying the relationship between these systems will be an important goal for future work.

Beyond its implications for models of speech processing, the compact yet hierarchically organized representations within PML also have practical significance. Emerging intracranial neural interfaces remain constrained by limited spatial sampling, making the choice of implantation site critical. Current speech neuroprostheses that target sensorimotor cortex primarily exploit low-level articulatory, phonemic, and motor-planning representations to decode intended speech (Silva et al., 2024; Card et al., 2024). The hierarchical organization identified here suggests that PML bridges these lower-level processes with progressively higher-level lexical and semantic information, providing access to a broad spectrum of linguistic information within a single compact cortical region. This makes PML a promising target for future speech neuroprosthetic applications, with the potential to extend beyond decoding speech articulation toward restoring communication in a broader range of speech and language disorders.

## Method

### MRI Dataset and preprocessing

We used previously collected natural speech perception data (LeBel et al. 2023) for this study. Briefly, 7 participants each listened to more than 6 hours of podcasts in a 3T MR scanner in separate sessions, while their BOLD activity was collected. Data used in this study included 27 10–15 minute stories from *The Moth Radio Hour*. One story was presented 5 times across 5 sessions of data collection and was used as the held out story for all modeling procedures.

Functional scans were collected using gradient-echo EPI with repetition time (TR) = 2.00 s, echo time (TE) = 30.8 ms, flip angle = 71°, multi-band factor (simultaneous multi-slice) = 2, voxel size = 2.6 mm × 2.6 mm × 2.6 mm (slice thickness = 2.6 mm). Anatomical data was collected using a T1-weighted multi-echo MPRAGE sequence on the same scanner with voxel size = 1 mm × 1 mm × 1 mm following the Freesurfer morphometry protocol.

Data preprocessing followed the pipeline provided by the same publication (LeBel et al. 2023). All data used were motion corrected and aligned to a template volume using the FMRIB Linear Image Registration Tool (FLIRT) from FMRIB Software Library (FSL). To account for low frequency voxel response drifts, data was then detrended using a second order Savitzky-Golay filter with a 120-second window. The first 50 seconds (25 TRs) and the last 20 seconds (10 TRs) of the data were trimmed to avoid artifacts from onset transients and poor detrending performance at the edges of the data. This included 10 seconds (5 TRs) of resting period and the beginning and end of each story. The data from each voxel were then demeaned and scaled to unit variance. Before any modeling and analyses, nuisance regression was conducted by estimating the effect of 6 motion parameters and 27 eye voxel BOLD signals on each cortical voxel using ridge regression. The model predictions were subtracted from the preprocessed signal and were used for all modeling.

### Localizer Tasks

We functionally defined premotor language areas (PML) in all subjects using the auditory and imagined speech tasks. Motor localizer tasks were performed to locate primary motor areas for mouth, hand, and feet, as well as premotor oculomotor areas. These tasks are included in the aforementioned dataset (LeBel et al. 2023). The motor areas are used to define intereffector areas for the somato-cognitive action network (SCAN; Gordon et al., 2023). We also localized the laryngeal motor cortex (LMC) in one subject (S02) to illustrate its relation with PML.

The auditory task data was collected in one 10-minute scan. Participants listened to 10 repeats of a 1-minute auditory stimulus containing 20 seconds of music, natural sounds, and speech. Explainable variance of voxel responses across repeats was calculated and used to localize posterior PML in individual subjects.

The motor localizers and the imagined speech tasks were collected in two identical 10-minute scans. Participants were cued to perform six different tasks in 20-second blocks: “mouth”, “hand”, “feet”, “saccade”, “speak”, and “rest”. In the “mouth” condition, participants were instructed to produce mouth movements (“blablabla”) without vocalization. In the “hand” condition, participants were instructed to drum their fingers on the bed without moving their arms. In the “feet” condition, participants were instructed to wiggle their toes without moving their legs. In the “saccade” condition, participants were instructed to saccade randomly between white dots presented on the screen. In the “speak” condition, participants were instructed to produce internal narratives about certain topics (e.g., “What did you have for breakfast today?”) without voicing or movements. Ordinary least squares regression models were fitted for each voxel across all six conditions, and the beta maps were used to functionally define the following ROIs in individual subjects: mouth, hand, and feet maps were used to define their corresponding primary motor areas (M1M, M1H, M1F); saccade maps were used to define oculomotor areas including supplementary eye fields (SEF), frontal eye fields (FEF), and premotor eye fields (PEF; or inferior frontal eye fields iFEF); speak maps were used to define language areas in the inferior frontal gyrus (IFG) as well as anterior PML.

For the LMC localizers, we adapted the individual effector localizer tasks from (Eichert et al., 2020). The participant was instructed to perform tongue retraction, lip protrusion, vowel production (“i-i-i-i-i-i”), and rest (breathing only) under controlled breathing instructions. Each trial consisted of 2 seconds of inhalation and 4 seconds of exhalation, with the movements in each condition carried out during the exhalation period. Ordinary least squares regression models were fitted for each voxel across all four conditions, and the beta map for vowel production was used to define the dLMC.

To test if PML is related to SCAN, we localized the intereffector regions by including voxels on the anterior surface of CS that do not belong to M1M, M1H, or M1F, as defined above. According to the functional connectivity sites in the SCAN paper, such voxels found between M1F and M1H were grouped into SCAN2, those between M1H and M1M into SCAN4. The inferior SCAN regions were farther away from our region of interest and were thus not localized for any analyses.

### Speech and semantic features

To understand how brain activity varied with different properties of natural speech, we extracted features from different levels of speech processing. We included two groups of speech features: (1) continuous levels of speech features from large speech models, and (2) a discrete set of features commonly used to map human speech processing in the brain.

First, continuous speech features were extracted from a large self-supervised speech model, WavLM-large (Chen et al., 2022). The stimulus waveforms were segmented using a sliding window of 16 seconds with a stride of 100 milliseconds, and the segments were then passed through the model. At all 25 layers of the model (including the CNN output), we extract the hidden states of the final token as the model’s representation for the window (Antonello et al., 2023).

In addition to continuous levels of speech features, we also used discrete levels, including spectral features, articulation (Mesgarani et al., 2014), word embeddings (English1000; Huth et al., 2016), and contextual embeddings (LLaMA-33B; Touvron et al., 2023). The spectral features were mel-band spectrograms with frequencies ranging from 0 Hz to 8 kHz with 256 windows (LeBel et al., 2021). Articulatory features were generated by mapping hand-labeled phonemes onto 14 features that describe the articulatory and acoustic properties of each phoneme (Chomsky & Halle, 1968; Mesgarani et al., 2014). For word embedding, we used the English1000 features, which capture lexical semantics in the stories (Huth et al., 2016). Finally, contextual semantic features were obtained from LLaMA-33B (Touvron et al., 2023), a large pretrained decoder-only Transformer language model. For each word in the transcribed stories, we extracted the hidden states of the last token from layer 18, using a dynamic context window to improve computational efficiency (Antonello et al., 2023).

For variance partitioning, we took features extracted from WavLM layer 9 and LLaMA layer 18. Although they mainly encode acoustic and semantic features, respectively, they are both affected by low-level signals such as word rate. To prevent the shared low-level signal from obscuring their unique contributions to our models, we regressed word rate out of each feature space before conducting the variance partitioning analyses.

### Voxelwise encoding models

To understand the selectivity for different speech features within PML, we modeled the neural responses as functions of each speech and language feature space using voxel-wise encoding models. To prepare the stimulus features for modeling, we temporally aligned them with MRI data and downsampled them with a 3-lobe Lanczos interpolation to match the temporal resolution of BOLD signals (0.5 Hz). After temporal alignment, features from each story are z-scored and concatenated together. The hemodynamic response functions for each voxel were modeled with a finite impulse response model, where separate weights were learned for each feature with multiple delays (2, 4, 6, and 8 seconds) (LeBel et al., 2023). This is done by stacking copies of the features, padded with zeros to the corresponding number of TRs. The BOLD signal from each voxel was then regressed onto this final stimulus matrix using bootstrapped ridge regression. The ridge regularization parameter for each voxel was selected independently using 50 iterations of cross-validation, with 50 random chunks of 40 TRs (∼20% of the total training data) used as the validation set. The performance of the encoding models was assessed with a held-out test story. Pearson’s correlation was calculated between model predictions and the actual BOLD responses for the test story and used as the final measure of model performance.

### Variance partitioning

To understand the unique contribution of acoustic and semantic features in different parts of PML, we estimated the unique variance explained ( ^2^) by each feature space. We first fit three encoding models for each voxel with: (A) speech features extracted from WavLM layer 9, (B) semantic features extracted from LLaMA layer 18, and (C) concatenated features from (A) and (B). We then calculated the variance explained in each voxel by each encoding model.

Using set-theory relations, we calculated the unique and shared variance explained by each feature space accordingly. More specifically, let *A* and *B* be the total variance explained by WavLM layer 9 features and LLaMA layer 18 features, respectively, and *A* ∪ *B* the total variance explained by the joint model (C). The shared variance explained by both feature spaces can thus be calculated as:

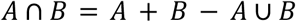

Then the unique variance explained by each feature was calculated as:

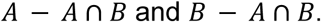

To create group maps that better display the general pattern across subjects, volumetric measures for each subject were first mapped to individual surface vertices using Pycortex’s mapper function. The resulting surface data were then mapped to Freesurfer’s template surface fsaverage using mri_surf2surf function (Fischl, 2012). The variance explained for each model were then averaged across subjects to produce the final map.

### Vertex Binning

To examine speech selectivity in PML along the anterior-posterior axis, we binned each vertex in PML by its position along the axis, enabling downstream analyses to quantify changes in measures as a function of the anatomical gradient. First, two vertices were identified on the most anterior and posterior borders of each localized PML. Following the methods introduced in (Popham et al., 2021), the shortest path along the cortical folds was identified between the two vertices and defined as the A-P axis for that ROI. This was done using a customized version of the Pycortex geodesic_path function to constrain the path to the PML vertices. Next, for each vertex *v_i_* in the ROI, we found its closest vertex *v_i_* on the path using the geodesic_distance function from Pycortex. The index *j* ∈ [1, 15] along the path was assigned as the position of *v_i_* along the A-P axis. Finally, the path was segmented into 15 equal-distance bins, and the vertices in PML were grouped into bins according to their positions along the axis.

To calculate model measures (prediction performance, variance explained) in each bin, volumetric measures for each subject were first mapped to individual surface vertices using Pycortex’s mapper function. Vertex-wise measures were averaged within each bin, then across subjects to obtain the final results. To show that PML representations vary along the anterior-posterior axis, we calculated the center of mass (CoM) for each feature space across bins (below).

### Task-State Functional Correlations (tsFC)

To investigate the functional similarity between PML and other areas, we calculated task-state functional correlations (tsFC) between the two areas. We used 6 sessions (over 5 hours) of story listening data and concatenated the responses. The beginning and end of each story scan were trimmed to eliminate ramping effects of the MRI signal (LeBel et al., 2023). For each seed voxel, we calculated its Pearson correlation with all voxels on the cortex.

For ROI-based tsFC, we first calculated the correlations seeded from all voxels within the ROI (e.g., AC, S1M, SCAN2), then averaged across the voxels to observe the overall effect. A two-way repeated-measures ANOVA was used to assess the effects of seed ROI, target ROI, and their interaction on tsFC. Following a significant seed ROI × target ROI, planned paired t-tests were performed to compare the matched seed-target connection with the two non-matched seed-target connections for each target. P-values from the six planned comparisons were corrected for multiple comparisons using false discovery rate (FDR) procedure (Benjamini & Hochberg, 1995) with threshold *p* < 0. 05.

When calculating the tsFC seeded from PML bins, we first calculated the correlations seeded from all voxels within PML. Next, the voxelwise results were interpolated and mapped to surface vertices using mappers from Pycortex with the “line_nearest” sampler, such that the value for each vertex is a weighted average of voxels that intersect with the line connecting the two corresponding vertices on the pial and white surfaces. The vertices were then binned by the vertex grouping introduced above, and the average tsFC was calculated for each bin. To show that they follow the anterior-posterior gradient, we calculated the center of mass (below) for tsFCs along the A-P line as the average of all bins weighted by their index.

### Center of Mass (CoM)

Encoding model performance and functional connectivity measures were evaluated in each bin *b* to reveal their spatial organization along the anterior-posterior axis. To quantify the profile for each measure *f* along the anterior-posterior gradient, we calculated the center of mass of its value *r* across all *B* bins as:

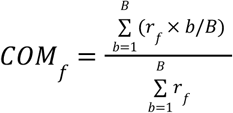

To demonstrate that PML representations vary along the anterior-posterior axis, we compared the CoMs for each feature set (all WavLM layers; 4 discrete speech features) with the bin indices. Pearson’s correlation was calculated between the CoMs in their corresponding order and the bin indices as the strength of gradient. To test if the correlations are significantly different from 0, we calculated Pearson’s correlation between randomly permuted CoMs and the bin indices. This was repeated 1000 times to establish the null distribution, which was used to calculate the p-value for each feature set. Multiple comparison correction was done with FDR (Benjamini & Hochberg, 1995) with a threshold of *p* < 0. 001.

### Significance testing

To test whether model measures (prediction performance and variance explained) and tsFC are significantly greater than 0, we conducted permutation tests for each measure. For model measures, block-wise permutations (block size 10 TRs) of model predictions were correlated with the real BOLD signal for the test story for each subject. For tsFC, seed voxel BOLD signal was permuted and correlated with target voxel signals. Then the final measures for each metric were obtained following the steps described above. This procedure was repeated 1000 times to establish the null distribution, which was used to calculate the p-value for each bin (for binned measures) or for each vertex (for whole cortex measures). Multiple comparison correction was done with FDR (Benjamini & Hochberg, 1995) with a threshold of *p* < 0. 001.

---

**Figure S1.**
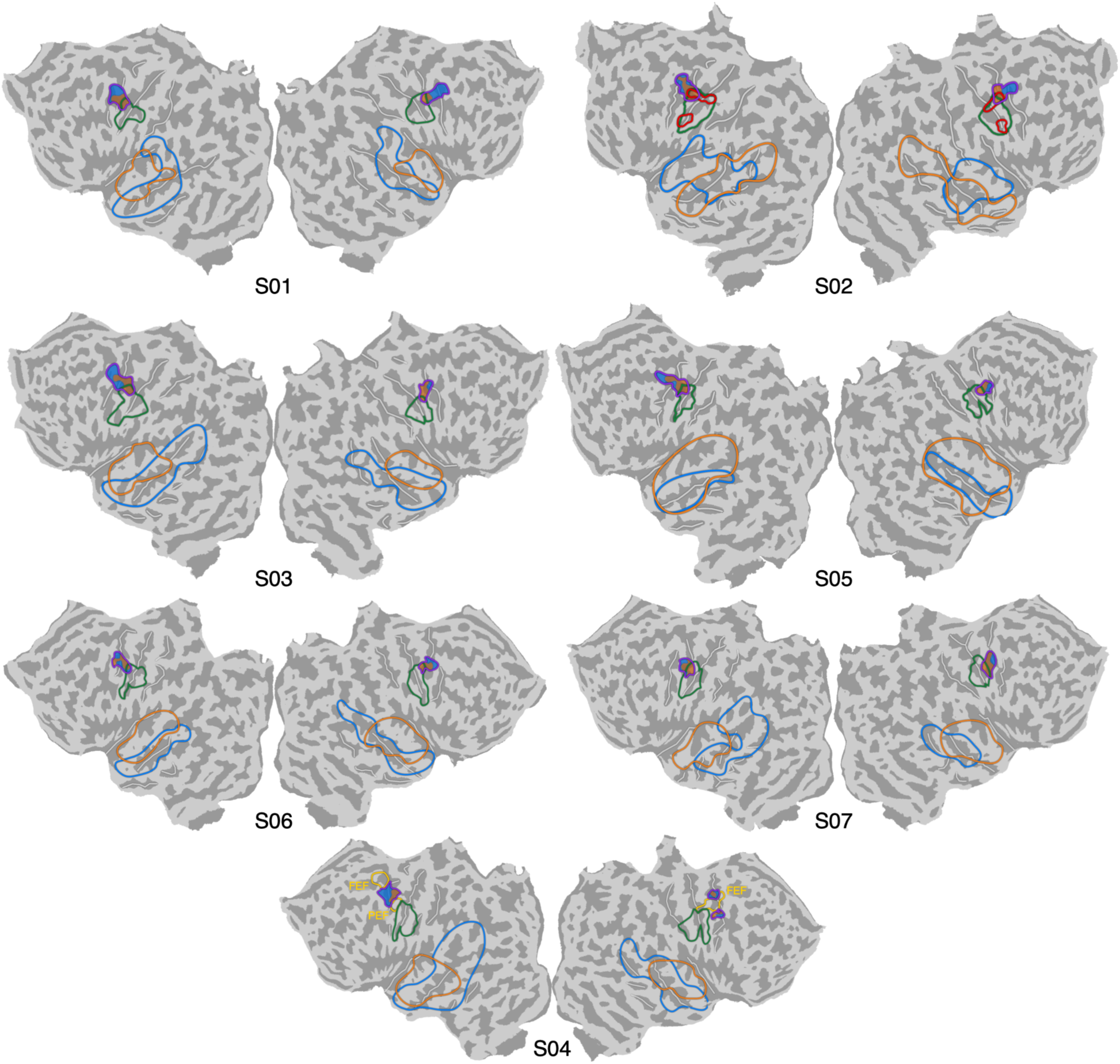
ROI localization for individual subjects. Similar to Figure 1 in the main text. Posterior PML is located on the crown of preCG, partially overlapping with the primary mouth motor cortex (M1M; green). Anterior PML is more variable: for some subjects, it is on the gyrus connecting preCG and MFG; for others, it passes through the anterior extension of the precentral sulcus. Inter-hemispheric differences were observed as well. S04 exhibited the rare organization of dual PML on the right hemisphere, where two unconnected regions sandwiching FEF (yellow) respond to both speech perception and production (Glasser et al., 2016). This subject was excluded from further analyses. Laryngeal motor cortex (LMC; red) is localized in S02 only. Purple: PML; orange: audition task activations; blue: imagined speech task activations; yellow: oculomotor areas (FEF, PEF). Six major sulci are marked in white: central sulcus, pre- and post-central sulci, Sylvian fissure, superior temporal sulcus, and inferior temporal sulcus.

**Figure S2.**
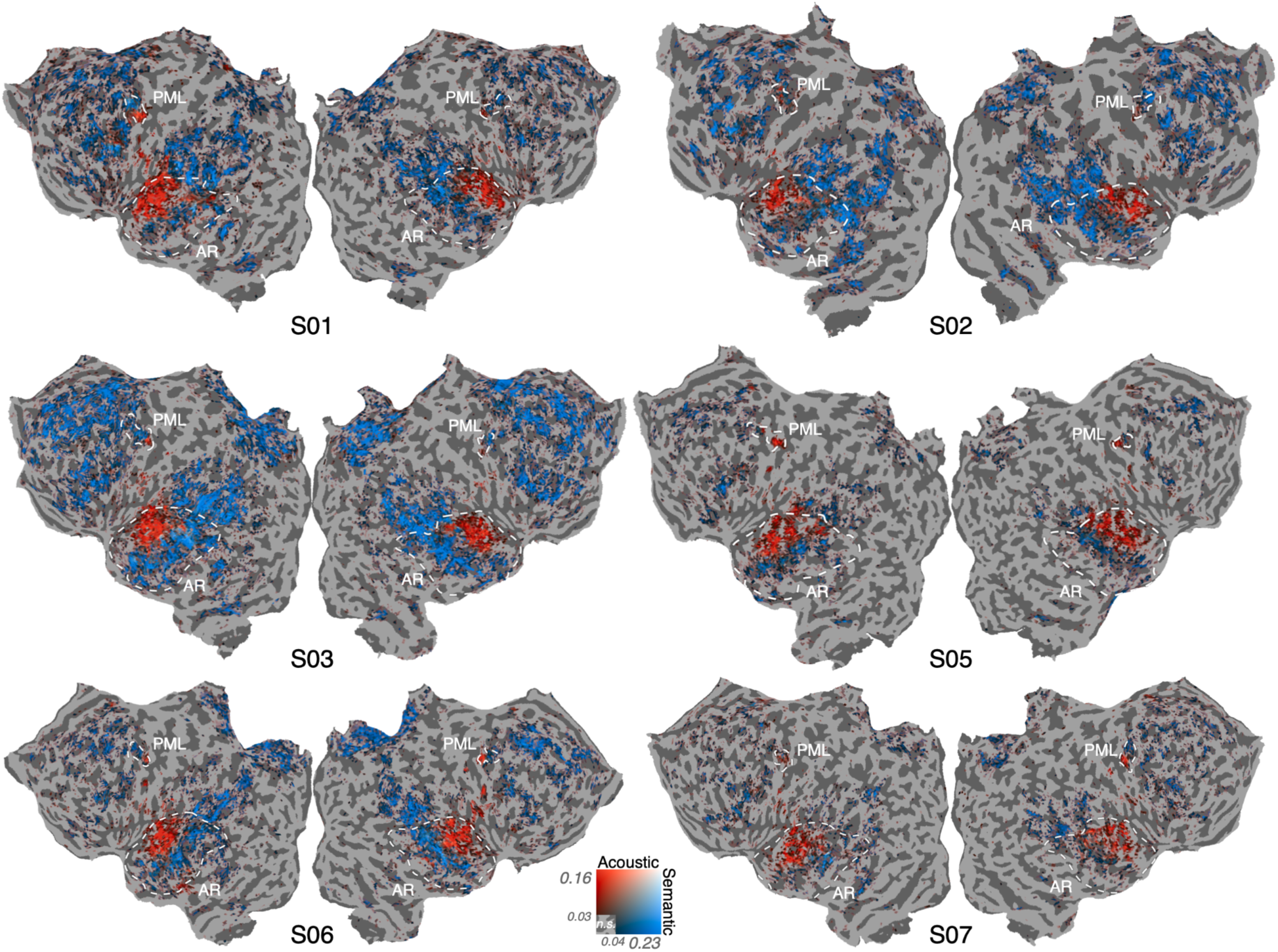
Variance partitioning for individual subjects. Similar to Figure 2D in the main text. WavLM layer 9 (acoustic, red) uniquely explains more variance in posterior PML and medial temporal auditory regions (AR), while LLaMA layer 18 (semantic, blue) uniquely explains more variance in anterior PML and lateral AR. Only voxels with significant amount of variance explained by either model are plotted in color here (*p* < 0. 001, permutation test, FDR corrected); in the main results all data are used to compute the average across subjects.

**Figure S3.**
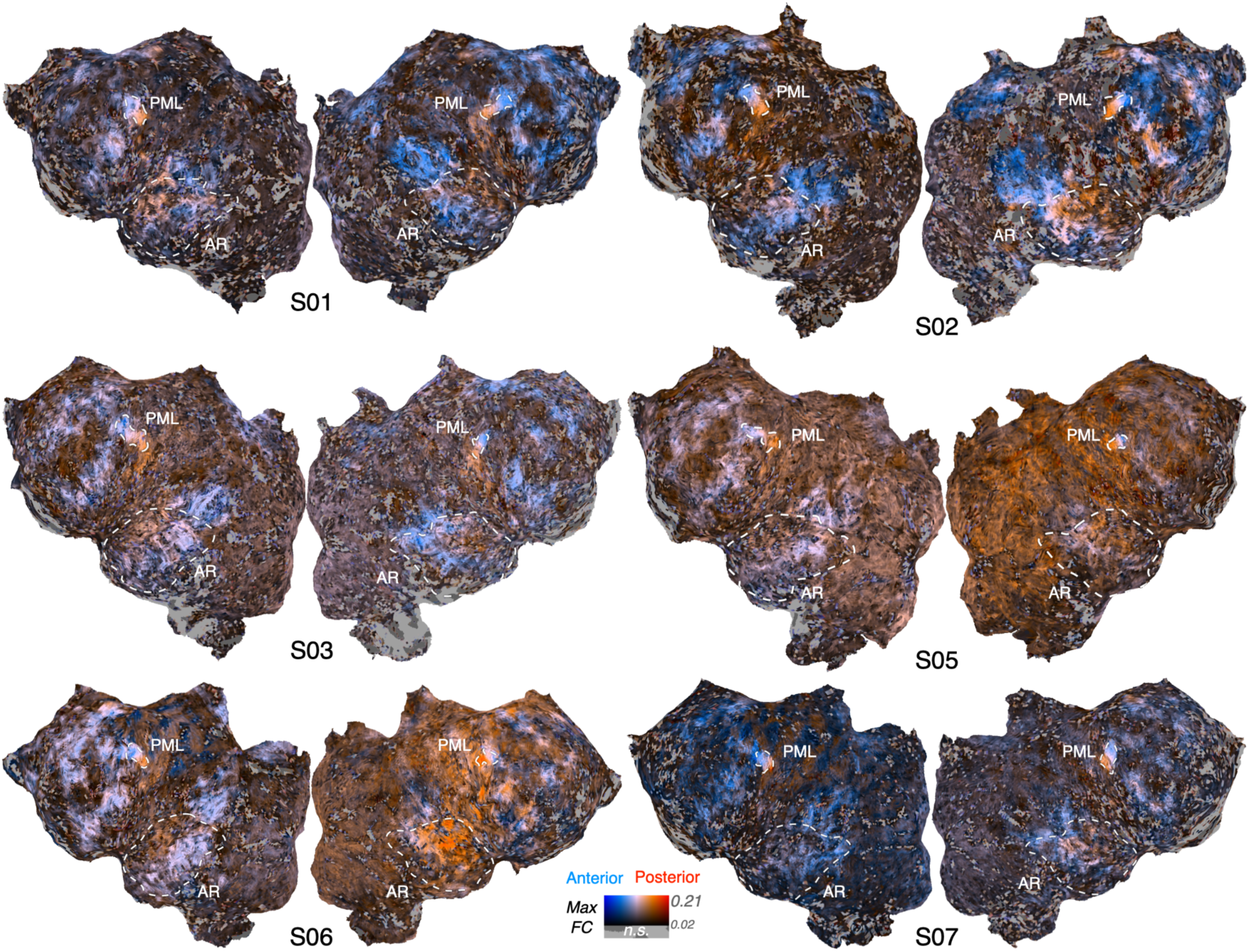
Task-state functional connectivity (tsFC) strength (hue) and preference to anterior (blue) or posterior (red) PML, for all cortical voxels in individual subjects. Similar to Figure 3A in the main text. The PML functional gradient is correlated with the known speech processing pathway in temporal AR. Voxels with significant tsFC in at least one bin are plotted in color (*p* < 0. 001, permutation test, FDR corrected); in the main results all data are used to compute the average across subjects.

**Figure S4.**
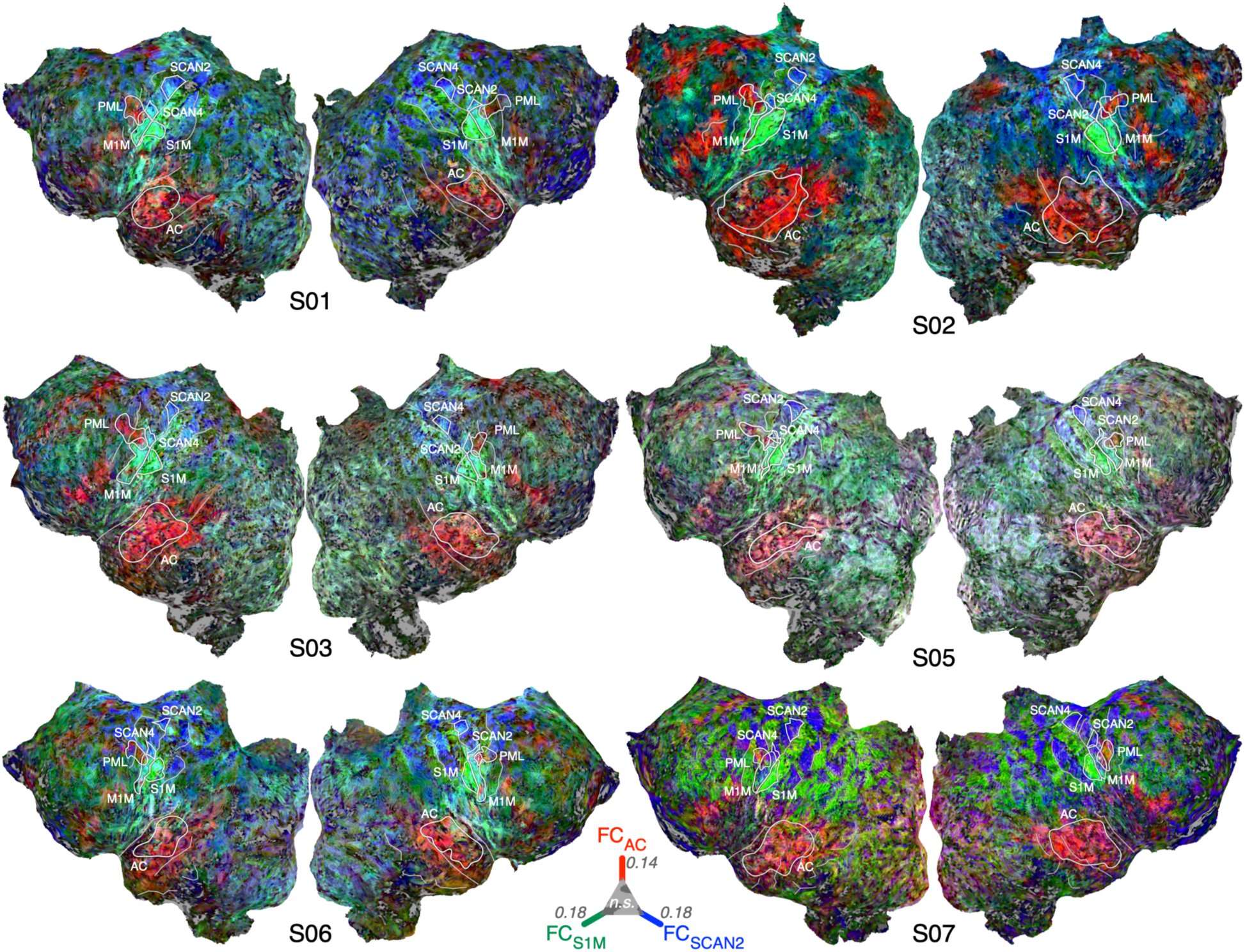
Task-state functional connectivity (tsFC) seeded from AC (red), S1M (green), and SCAN2 (blue). Similar to Figure 3B in the main text. The strongest correlations are between AC-PML, S1M-M1M, and SCAN2-SCAN4. Voxels with significant tsFC from at least one seed ROI are plotted in color (*p* < 0. 001, permutation test, FDR corrected); in the main results all data are used to compute the average across subjects.

## References

Antonello, R., Vaidya, A., & Huth, A. (2023). Scaling laws for language encoding models in fMRI. Advances in Neural Information Processing Systems, 36, 21895–21907.

Ardila, A. (2020). Supplementary motor area aphasia revisited. Journal of Neurolinguistics, 54, 100888. 10.1016/j.jneuroling.2020.100888

Axer, H., Klingner, C. M., & Prescher, A. (2013). Fiber anatomy of dorsal and ventral language streams. Brain and Language, 127(2), 192–204. 10.1016/j.bandl.2012.04.015

Benjamini, Y., & Hochberg, Y. (1995). Controlling the False Discovery Rate: A Practical and Powerful Approach to Multiple Testing. Journal of the Royal Statistical Society: Series B (Methodological), 57(1), 289–300. 10.1111/j.2517-6161.1995.tb02031.x

Callan, D., Callan, A., Gamez, M., Sato, M., & Kawato, M. (2010). Premotor cortex mediates perceptual performance. NeuroImage, 51(2), 844–858. 10.1016/j.neuroimage.2010.02.027

Callan, D., Jones, J. A., Callan, A. M., & Akahane-Yamada, R. (2004). Phonetic perceptual identification by native- and second-language speakers differentially activates brain regions involved with acoustic phonetic processing and those involved with articulatory–auditory/orosensory internal models. NeuroImage, 22(3), 1182–1194. 10.1016/j.neuroimage.2004.03.006

Card, N. S., Wairagkar, M., Iacobacci, C., Hou, X., Singer-Clark, T., Willett, F. R., Kunz, E. M., Fan, C., Nia, M. V., Deo, D. R., Srinivasan, A., Choi, E. Y., Glasser, M. F., Hochberg, L. R., Henderson, J. M., Shahlaie, K., Stavisky, S. D., & Brandman, D. M. (2024). An Accurate and Rapidly Calibrating Speech Neuroprosthesis. New England Journal of Medicine, 391(7), 609–618. 10.1056/NEJMoa2314132

Chen, S., Wang, C., Chen, Z., Wu, Y., Liu, S., Chen, Z., Li, J., Kanda, N., Yoshioka, T., Xiao, X., Wu, J., Zhou, L., Ren, S., Qian, Y., Qian, Y., Wu, J., Zeng, M., Yu, X., & Wei, F. (2022). WavLM: Large-Scale Self-Supervised Pre-Training for Full Stack Speech Processing. IEEE Journal of Selected Topics in Signal Processing, 16(6), 1505–1518. 10.1109/JSTSP.2022.3188113

Cheung, C., Hamilton, L. S., Johnson, K., & Chang, E. F. (2016). The auditory representation of speech sounds in human motor cortex. eLife, 5, e12577. 10.7554/eLife.12577

Cho, C. J., Wu, P., Prabhune, T. S., Agarwal, D., & Anumanchipalli, G. K. (2024). Coding Speech Through Vocal Tract Kinematics. IEEE Journal of Selected Topics in Signal Processing, 18(8), 1427–1440. 10.1109/JSTSP.2024.3497655

Chomsky, N., & Halle, M. (1968). *The sound pattern of English*.

Correia, J. M., Caballero-Gaudes, C., Guediche, S., & Carreiras, M. (2020). Phonatory and articulatory representations of speech production in cortical and subcortical fMRI responses. Scientific Reports, 10(1), 4529. 10.1038/s41598-020-61435-y

D’Ausilio, A., Bufalari, I., Salmas, P., Busan, P., & Fadiga, L. (2011). Vocal pitch discrimination in the motor system. Brain and Language, 118(1), 9–14. 10.1016/j.bandl.2011.02.007

Dichter, B. K., Breshears, J. D., Leonard, M. K., & Chang, E. F. (2018). The Control of Vocal Pitch in Human Laryngeal Motor Cortex. Cell, 174(1), 21–31.e9. 10.1016/j.cell.2018.05.016

Dick, A. S., Bernal, B., & Tremblay, P. (2014). The Language Connectome: New Pathways, New Concepts. The Neuroscientist, 20(5), 453–467. 10.1177/1073858413513502

Du, Y., Buchsbaum, B. R., Grady, C. L., & Alain, C. (2014). Noise differentially impacts phoneme representations in the auditory and speech motor systems. Proceedings of the National Academy of Sciences, 111(19), 7126–7131. 10.1073/pnas.1318738111

Eichert, N., Papp, D., Mars, R. B., & Watkins, K. E. (2020). Mapping Human Laryngeal Motor Cortex during Vocalization. Cerebral Cortex, 30(12), 6254–6269. 10.1093/cercor/bhaa182

Fedorenko, E., Hsieh, P.-J., Nieto-Castañón, A., Whitfield-Gabrieli, S., & Kanwisher, N. (2010). New Method for fMRI Investigations of Language: Defining ROIs Functionally in Individual Subjects. Journal of Neurophysiology, 104(2), 1177–1194. 10.1152/jn.00032.2010

Fedorenko, E., Ivanova, A. A., & Regev, T. I. (2024). The language network as a natural kind within the broader landscape of the human brain. Nature Reviews Neuroscience, 25(5), 289–312. 10.1038/s41583-024-00802-4

Fischl, B. (2012). FreeSurfer. NeuroImage, 20 YEARS OF fMRI, 62(2), 774–781. 10.1016/j.neuroimage.2012.01.021

Glasser, M. F., Coalson, T. S., Robinson, E. C., Hacker, C. D., Harwell, J., Yacoub, E., Ugurbil, K., Andersson, J., Beckmann, C. F., Jenkinson, M., Smith, S. M., & Van Essen, D. C. (2016). A multi-modal parcellation of human cerebral cortex. Nature, 536(7615), 171–178. 10.1038/nature18933

Gordon, E. M., Chauvin, R. J., Van, A. N., Rajesh, A., Nielsen, A., Newbold, D. J., Lynch, C. J., Seider, N. A., Krimmel, S. R., Scheidter, K. M., Monk, J., Miller, R. L., Metoki, A., Montez, D. F., Zheng, A., Elbau, I., Madison, T., Nishino, T., Myers, M. J., … Dosenbach, N. U. F. (2023). A somato-cognitive action network alternates with effector regions in motor cortex. Nature, 617(7960), 351–359. 10.1038/s41586-023-05964-2

Hamilton, L. S., Edwards, E., & Chang, E. F. (2018). A Spatial Map of Onset and Sustained Responses to Speech in the Human Superior Temporal Gyrus. Current Biology, 28(12), 1860–1871.e4. 10.1016/j.cub.2018.04.033

Heer, W. A. de, Huth, A. G., Griffiths, T. L., Gallant, J. L., & Theunissen, F. E. (2017). The Hierarchical Cortical Organization of Human Speech Processing. Journal of Neuroscience, 37(27), 6539–6557. 10.1523/JNEUROSCI.3267-16.2017

Hertrich, I., Dietrich, S., & Ackermann, H. (2016). The role of the supplementary motor area for speech and language processing. Neuroscience & Biobehavioral Reviews, 68, 602–610. 10.1016/j.neubiorev.2016.06.030

Hickok, G., & Poeppel, D. (2004). Dorsal and ventral streams: A framework for understanding aspects of the functional anatomy of language. *Cognition*, Towards a New Functional Anatomy of Language, 92(1), 67–99. 10.1016/j.cognition.2003.10.011

Hickok, G., Venezia, J., & Teghipco, A. (2023). Beyond Broca: Neural architecture and evolution of a dual motor speech coordination system. Brain, 146(5), 1775–1790. 10.1093/brain/awac454

Higashiyama, Y., Hamada, T., Saito, A., Morihara, K., Okamoto, M., Kimura, K., Joki, H., Kishida, H., Doi, H., Ueda, N., Takeuchi, H., & Tanaka, F. (2021). Neural mechanisms of foreign accent syndrome: Lesion and network analysis. NeuroImage: Clinical, 31, 102760. 10.1016/j.nicl.2021.102760

Hopf, A. (1956). Über die Verteilung myeloarchitektonischer Merkmale in der Stirnhirnrinde beim Menschen. J Hirnforsch, 2(4), 311–333.

Huth, A. G., de Heer, W. A., Griffiths, T. L., Theunissen, F. E., & Gallant, J. L. (2016). Natural speech reveals the semantic maps that tile human cerebral cortex. Nature, 532(7600), 453–458. 10.1038/nature17637

LeBel, A., Jain, S., & Huth, A. G. (2021). Voxelwise Encoding Models Show That Cerebellar Language Representations Are Highly Conceptual. Journal of Neuroscience, 41(50), 10341–10355. 10.1523/JNEUROSCI.0118-21.2021

LeBel, A., Wagner, L., Jain, S., Adhikari-Desai, A., Gupta, B., Morgenthal, A., Tang, J., Xu, L., & Huth, A. G. (2023). A natural language fMRI dataset for voxelwise encoding models. Scientific Data, 10(1), 555. 10.1038/s41597-023-02437-z

Levy, D., Greicius, Q., Wang, C., Ko, E., Xu, D., Andrews, J., & Chang, E. F. (2025). Role for Left Dorsomedial Prefrontal Cortex in Self-Generated, but not Externally Cued, Language Production. Neurobiology of Language, 6, nol_a_00166. 10.1162/nol_a_00166

Lu, J., Zhao, Z., Zhang, J., Wu, B., Zhu, Y., Chang, E. F., Wu, J., Duffau, H., & Berger, M. S. (2021). Functional maps of direct electrical stimulation-induced speech arrest and anomia: A multicentre retrospective study. Brain, 144(8), 2541–2553. 10.1093/brain/awab125

Meister, I. G., Wilson, S. M., Deblieck, C., Wu, A. D., & Iacoboni, M. (2007). The Essential Role of Premotor Cortex in Speech Perception. Current Biology, 17(19), 1692–1696. 10.1016/j.cub.2007.08.064

Mesgarani, N., Cheung, C., Johnson, K., & Chang, E. F. (2014). Phonetic Feature Encoding in Human Superior Temporal Gyrus. Science, 343(6174), 1006–1010. 10.1126/science.1245994

Mohamed, A., Lee, H., Borgholt, L., Havtorn, J. D., Edin, J., Igel, C., Kirchhoff, K., Li, S.-W., Livescu, K., Maaløe, L., Sainath, T. N., & Watanabe, S. (2022). Self-Supervised Speech Representation Learning: A Review. IEEE Journal of Selected Topics in Signal Processing, 16(6), 1179–1210. 10.1109/JSTSP.2022.3207050

Pasad, A., Shi, B., & Livescu, K. (2023). Comparative Layer-Wise Analysis of Self-Supervised Speech Models. ICASSP 2023 - 2023 IEEE International Conference on Acoustics, Speech and Signal Processing (ICASSP), 1–5. 10.1109/ICASSP49357.2023.10096149

Paus, T., Perry, D. W., Zatorre, R. J., Worsley, K. J., & Evans, A. C. (1996). Modulation of Cerebral Blood Flow in the Human Auditory Cortex During Speech: Role of Motor-to-sensory Discharges. European Journal of Neuroscience, 8(11), 2236–2246. 10.1111/j.1460-9568.1996.tb01187.x

Popham, S. F., Huth, A. G., Bilenko, N. Y., Deniz, F., Gao, J. S., Nunez-Elizalde, A. O., & Gallant, J. L. (2021). Visual and linguistic semantic representations are aligned at the border of human visual cortex. Nature Neuroscience, 24(11), 1628–1636. 10.1038/s41593-021-00921-6

Price, C. J. (2012). A review and synthesis of the first 20 years of PET and fMRI studies of heard speech, spoken language and reading. NeuroImage, 20 *YEARS OF fMRI*, *62*(2), 816–847. 10.1016/j.neuroimage.2012.04.062

Pulvermüller, F., Huss, M., Kherif, F., Moscoso del Prado Martin, F., Hauk, O., & Shtyrov, Y. (2006). Motor cortex maps articulatory features of speech sounds. Proceedings of the National Academy of Sciences, 103(20), 7865–7870. 10.1073/pnas.0509989103

Romanski, L. M., Tian, B., Fritz, J., Mishkin, M., Goldman-Rakic, P. S., & Rauschecker, J. P. (1999). Dual streams of auditory afferents target multiple domains in the primate prefrontal cortex. Nature Neuroscience, 2(12), 1131–1136. 10.1038/16056

Sammler, D., Grosbras, M.-H., Anwander, A., Bestelmeyer, P. E. G., & Belin, P. (2015). Dorsal and Ventral Pathways for Prosody. Current Biology, 25(23), 3079–3085. 10.1016/j.cub.2015.10.009

Schmahmann, J. D., Pandya, D. N., Wang, R., Dai, G., D’Arceuil, H. E., de Crespigny, A. J., & Wedeen, V. J. (2007). Association fibre pathways of the brain: Parallel observations from diffusion spectrum imaging and autoradiography. Brain, 130(3), 630–653. 10.1093/brain/awl359

Shekari, E., & Nozari, N. (2023). A narrative review of the anatomy and function of the white matter tracts in language production and comprehension. Frontiers in Human Neuroscience, 17. 10.3389/fnhum.2023.1139292

Silva, A. B., Littlejohn, K. T., Liu, J. R., Moses, D. A., & Chang, E. F. (2024). The speech neuroprosthesis. Nature Reviews Neuroscience, 25(7), 473–492. 10.1038/s41583-024-00819-9

Silva, A. B., Liu, J. R., Zhao, L., Levy, D. F., Scott, T. L., & Chang, E. F. (2022). A Neurosurgical Functional Dissection of the Middle Precentral Gyrus during Speech Production. Journal of Neuroscience, 42(45), 8416–8426. 10.1523/JNEUROSCI.1614-22.2022

Tate, M. C., Herbet, G., Moritz-Gasser, S., Tate, J. E., & Duffau, H. (2014). Probabilistic map of critical functional regions of the human cerebral cortex: Broca’s area revisited. Brain, 137(10), 2773–2782. 10.1093/brain/awu168

Touvron, H., Lavril, T., Izacard, G., Martinet, X., Lachaux, M.-A., Lacroix, T., Rozière, B., Goyal, N., Hambro, E., Azhar, F., Rodriguez, A., Joulin, A., Grave, E., & Lample, G. (2023). *LLaMA: Open and Efficient Foundation Language Models* (arXiv:2302.13971). arXiv. 10.48550/arXiv.2302.13971

Tremblay, P., & Small, S. L. (2011). On the context-dependent nature of the contribution of the ventral premotor cortex to speech perception. NeuroImage, 57(4), 1561–1571. 10.1016/j.neuroimage.2011.05.067

Vaidya, A. R., Jain, S., & Huth, A. G. (2022). *Self-supervised models of audio effectively explain human cortical responses to speech* (arXiv:2205.14252). arXiv. 10.48550/arXiv.2205.14252

Venezia, J. H., Richards, V. M., & Hickok, G. (2021). Speech-Driven Spectrotemporal Receptive Fields Beyond the Auditory Cortex. Hearing Research, 408, 108307. 10.1016/j.heares.2021.108307

Vigneau, M., Beaucousin, V., Hervé, P. Y., Duffau, H., Crivello, F., Houdé, O., Mazoyer, B., & Tzourio-Mazoyer, N. (2006). Meta-analyzing left hemisphere language areas: Phonology, semantics, and sentence processing. NeuroImage, 30(4), 1414–1432. 10.1016/j.neuroimage.2005.11.002

Wilson, S. M., & Iacoboni, M. (2006). Neural responses to non-native phonemes varying in producibility: Evidence for the sensorimotor nature of speech perception. NeuroImage, 33(1), 316–325. 10.1016/j.neuroimage.2006.05.032

Wilson, S. M., Saygin, A. P., Sereno, M. I., & Iacoboni, M. (2004). Listening to speech activates motor areas involved in speech production. Nature Neuroscience, 7(7), 701–702. 10.1038/nn1263

Wong, P. C. M., Uppunda, A. K., Parrish, T. B., & Dhar, S. (2008). Cortical Mechanisms of Speech Perception in Noise. Journal of Speech, Language, and Hearing Research, 51(4), 1026–1041. 10.1044/1092-4388(2008/075)

Wu, G., Wang, X., Lyu, B., & Du, Y. (2026). Area 55b forms a cortical transition zone for sensorimotor-semantic transformation in human language (Rs. 3.rs-9934100). Research Square. 10.21203/rs.3.rs-9934100/v1

